# A primary culture method for the easy, efficient and effective acquisition of oligodendrocyte lineage cells

**DOI:** 10.1101/2023.11.27.568774

**Authors:** Hanki Kim, Bum Jun Kim, Seungyon Koh, Hyo Jin Cho, Xuelian Jin, Byung Gon Kim, Jun Young Choi

**Author notes:** Correspondence: Jun Young Choi M.D., Ph.D., Departments of Brain Science and Neurology, Ajou University School of Medicine, Suwon, 16499, Korea. **Conflict of interest statement:** The authors declare no conflicts of interest.

## Abstract

Oligodendrocytes (OL) are myelin forming glial cells in the central nervous system. *In vitro* primary OL culture models provide the benefit of a more readily controlled environment facilitating the evaluation of diverse OL stages and convoluted dynamics. Conventional methods of primary OL culture do exist, but their performance in terms of efficiency and simplicity has room for improvement. We present a novel method of primary OL culture, namely the E3 (Easy, efficient, and effective) method, which greatly improves cell yield and reduces time required to oligodendrocyte progenitor cell (OPC) acquisition and maturation into OLs. We also provide optimal media compositions for augmentation of OPC proliferation and a more robust maturation into myelin forming OLs. *In vitro* characteristics of the OL lineage discovered during the development of the E3 method present implications for further research on OL physiology and pathophysiology.

## 1. Introduction

Oligodendrocytes (OL) are the myelin-forming cells of the central nervous system (CNS), facilitating fast neuronal signal transmission by enabling saltatory conduction, as well as providing metabolic support to the neurons they enwrap (Bradl & Lassmann, 2010; Philips & Rothstein, 2017). OL death and demyelination causes neurological deficits, and insufficient remyelination eventually leads to axonal degeneration (Irvine & Blakemore, 2008; Simkins et al., 2021). Demyelination is implicated in various neurological disorders, the most renowned being multiple sclerosis (MS) and ischemic white matter injury (Duncan & Radcliff, 2016). Numerous efforts have been and are being made to study the development and characteristics of OLs, their role in the pathogenesis of demyelinating disorders, and methods to promote remyelination (Deshmukh et al., 2013). The complexity of the OL lineage makes it difficult to study their dynamics solely in an *in vivo* system where the diversity of OL stages spanning from oligodendrocyte progenitor cells (OPCs) to mature OLs, signals and interactions with other cell types may complicate the analysis of a single component, calling for the need of an *in vitro* model to perform detailed experiments.

*In vitro* OL culture models provide the advantage of a strictly regulated environment where OL lineage cells are isolated from other elements of the brain, at the cost of structure, signals and interactions that are preserved in an *in vivo* system. While a few established OL cell lines for culture do exist, they fail to fully recapitulate OL morphology and function, and do not span the whole OL lineage (Buntinx et al., 2003; Pereira et al., 2011). Thus, a large proportion of *in vitro* research on OLs, developmental myelination and de/remyelination in disease states has been conducted with primary-cultured OLs. Various methods to isolate OPCs from rodent brains are present, the most widely used being the shaking method which utilizes the different surface adhesion capacities of glial cells to shake off OPCs from a stratified monolayer of astrocytes in a mixed glial culture (McCarthy & de Vellis, 1980; O’Meara et al., 2011). Methods that exploit antigen-antibody reactions such as immunopanning or immunomagnetic cell separation also exist (Dugas & Emery, 2013b; Emery & Dugas, 2013; Weil et al., 2019). However, such methods tend to be costly, labor-intensive, and may also depend on the experimenter’s technique to accomplish desired outcomes. Recently, novel, more simple methods for OL culture have been devised, making use of OL-favoring growth factors or density gradient centrifugation to isolate OPCs from other brain cells (Nihonmatsu-Kikuchi et al., 2021; Yoshida et al., 2020). However, these methods have their own weaknesses in terms of cell yield or the purity of obtained cell population.

In this study, we present a new method of primary OL culture, which we term the E3 (easy, efficient and effective) method. The E3 method is a compilation of existing new and old methods for OPC isolation, along with the use of optimal media compositions for the proliferation and differentiation of isolated OPCs and OLs. The E3 method attained a high purity of OPCs comparable to conventional shaking methods, without overcomplicating the simple procedure which is the greatest advantage of the novel ones. Also, we present optimal media compositions for the achievement of robust OPC proliferation and differentiation into complex mature oligodendrocytes.

## 2. Materials and methods

### 2.1. OPC isolation

A detailed protocol containing information on reagents and equipment, and instructions for the full culture procedure, is available in the Supporting Information section. P1 rat pups were briefly placed on ice for anesthetization and decapitated. The heads were collected in a 50mL conical tube containing ice-cold Hank’s Balanced Salt Solution (HBSS), transferred to another tube containing ice-cold 70% ethanol, and were transferred again to a new tube with ice-cold HBSS. The cerebral cortices were dissected, and after the meninges had been removed, the hemispheres were moved to a 15mL conical tube containing 200_μ_Ls of HBSS, one brain (two hemispheres) per tube. 1mL of Accumax^TM^ (Merck Millipore, #SCR006) was added to each tube, and the brains were briefly triturated 10 times with a 1mL pipette tip. 4mLs of dissection media which consists of Hibernate-A medium (Thermo Fisher Scientific, A1247501), 1% GlutaMAX^TM^ (Thermo Fisher Scientific, #35050-061), 1% penicillin/streptomycin (Cytiva, #SV30010), plus 200IUs of Papain (Worthington, #LS003126), were added to each tube. The tissue was dissociated for 30 min at 30°C. The dissociated cell suspension was then centrifuged at 200g, 5 min, room temperature (RT), the supernatant was discarded, and the pellet was resuspended in 1mL of dissection media +12% Optiprep^TM^ (Sigma Aldrich, #D1556). To avoid the limitation of tissue processable associated with traditional density gradient centrifugation methods, we adopted the principle of differential centrifugation, a method utilizing different sedimentation speed of particles and commonly used for cellular organelle fractionation or exosome isolation (Claude, 1946; Livshits et al., 2016). The cell suspension was gently triturated 20 times with a 10_μ_L pipette tip attached to a 1mL tip. 3mLs of dissection media+12% Optiprep^TM^ was added, and the total 4mL cell suspension was centrifuged at 200g, 15 min, RT. The 4mL supernatant was transferred to a different 15mL tube, diluted with 4mLs of dissection media, and the resulting 8mL, 6% Optiprep^TM^ cell suspension was centrifuged at 200g, 15 min, RT. The final resulting pellet was resuspended in PBS, and cells were counted and seeded on poly-D-lysine (PDL, Sigma Aldrich, #P6407) coated culture surfaces (working concentration: 0.1mg/mL diluted in distilled water (DW), coated for 1 hour, washed with DW three times and air dried) at a density of 1*10^4^cells/cm^2^ in OPC proliferation media (**Table 1a**).

**Table 1.**
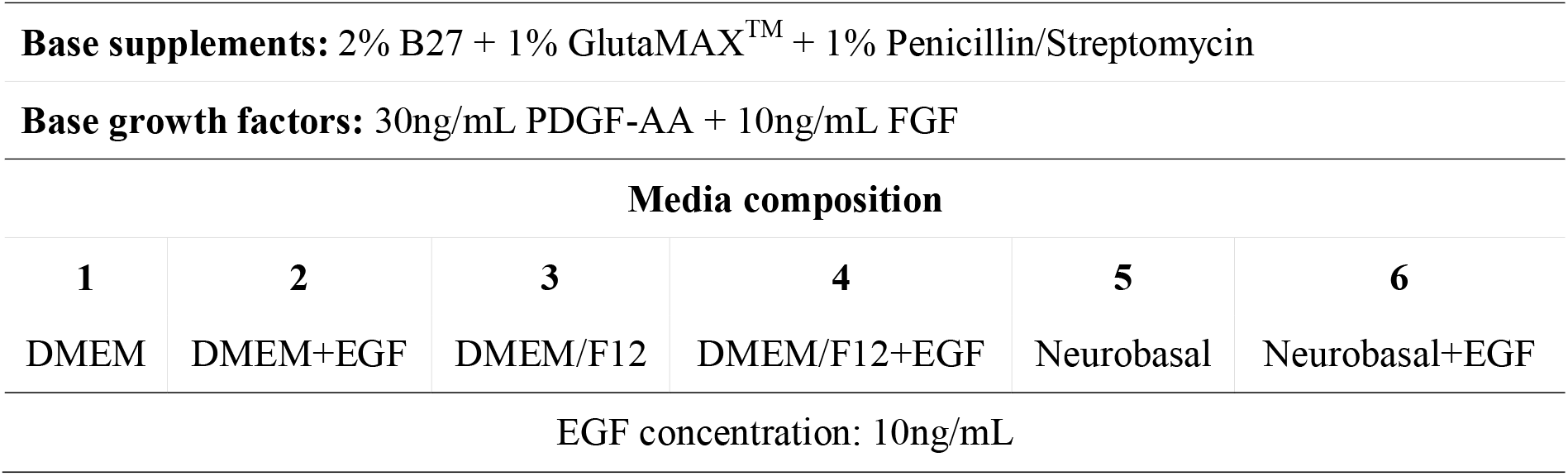
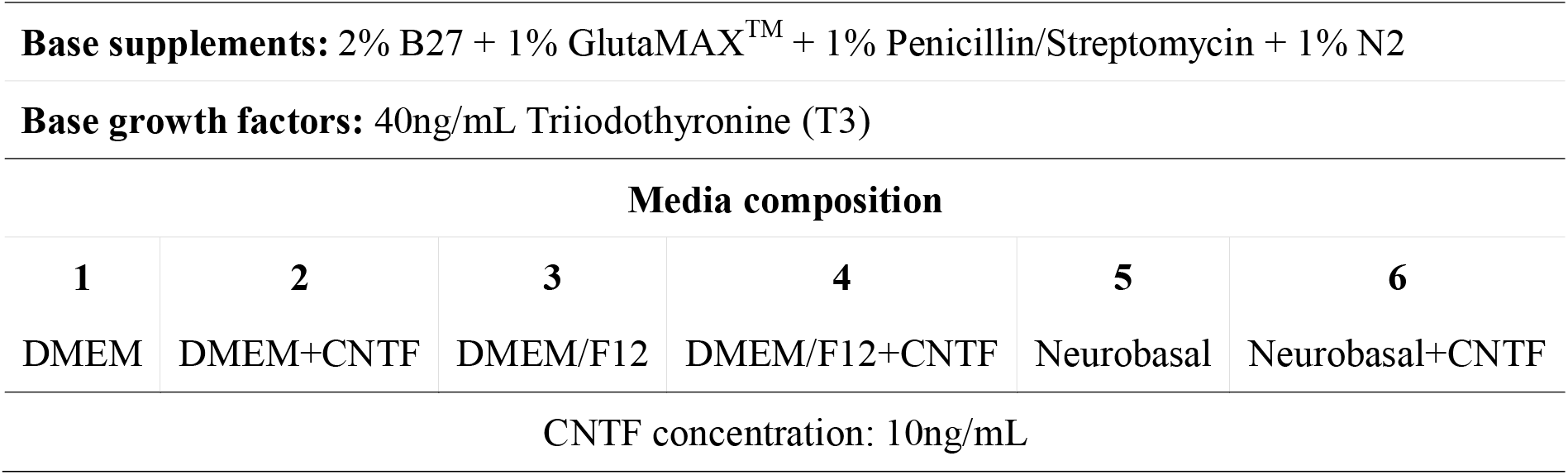
Media compositionss.

| Media composition |  |  |  |  |  |
| --- | --- | --- | --- | --- | --- |
| 1 | 2 | 3 | 4 | 5 | 6 |
| DMEM | DMEM+EGF | DMEM/F12 | DMEM/F12+EGF | Neurobasal | Neurobasal+EGF |
| EGF concentration: 10ng/mL |  |  |  |  |  |
| 1 | 2 | 3 | 4 | 5 | 6 |
| DMEM | DMEM+CNTF | DMEM/F12 | DMEM/F12+CNTF | Neurobasal | Neurobasal+CNTF |
| CNTF concentration: 10ng/mL |  |  |  |  |  |

Conventional mixed glial culture based OPC isolation methods were also performed (Choi et al., 2017; Choi et al., 2023; O’Meara et al., 2011). The sacrifice of pups, dissection, and dissociation was done as above, and the pellet was resuspended/triturated in 4mLs of Dulbecco’s modified eagle medium/Ham’s F12 (DMEM/F12, Thermo Fisher Scientific, #11320033), 10% fetal bovine serum (FBS), 1% GlutaMAX^TM^, 1% penicillin/streptomycin and centrifuged at 200g, 5 min. The pellet was resuspended in DMEM/F12, 10% (FBS), 1% GlutaMAX^TM^, 1% penicillin/streptomycin and seeded on PDL coated T-75 flasks, one brain/flask.

All animal experiments were reviewed and approved by the Institutional Animal Care and Use Committee of Ajou University School of Medicine (IACUC number; 2023-0021) and complied with the National Institutes of Health (NIH) Guide for the Care and Use of Laboratory Animals. Postnatal day 1 (P1) Sprague-Dawley rats were purchased from Orient Bio Korea, Inc.

### 2.2. OPC proliferation, passaging, and differentiation

Isolated OPCs were proliferated in OPC proliferation media of varying conditions (**Table 1a**) for 5-6 days, with a full media change at day 3 of proliferation. For further passaging and differentiation, OPC proliferation media, condition 4 of **Table 1a** was chosen. After the proliferation period, the media was aspirated and 5mL/T75 flask of Accutase® (Thermo Fisher Scientific, #A11105-01) was added, and the flask(s) were incubated in a 37°C, 5% CO_2_ incubator for 5 min. The flask(s) were gently tapped to detach cells, and the cell suspension was transferred to a 15mL tube and centrifuged at 200g, 3min, room temperature (RT). The resulting pellet was resuspended in PBS, counted, and seeded on appropriate culture surfaces at a density of 1*10^4^ cells/cm^2^ and proliferated for 2 days. For further differentiation, passaged OPCs cultured in condition 3 of **Table 1a** were chosen. A full media exchange to OL differentiation media (**Table 1b**) was performed, and cells were differentiated for 2 or 4 days.

### 2.3. Oligodendrocyte/Neuron co-culture

Primary neurons were obtained from P1 rat cortices by the following method. From the OPC isolation method in 2.2, cells pelleted from the first centrifuge in dissection media+12% Optiprep^TM^ were seeded on culture surfaces at a density of 5*10^4^ cells/cm^2^, in a media composition of Neurobasal medium (NBM, Thermo Fisher Scientific, #21103049), 2% B27 Supplement (Thermo Fisher Scientific, #17504-044), 1% N2 supplement (Thermo Fisher Scientific, #17502048), 1% GlutaMAX^TM^, 1% penicillin/streptomycin. The cells were maintained for 14 days, with a half media exchange every 3 days, which yielded a primary neuron culture. On the 14^th^ day, the media was exchanged to OL differentiation media, condition 5 of **Table 1b** and passaged OPCs were seeded onto the neurons at a density of 3*10^4^ cells/cm^2^. The co-culture was maintained for 4 days, with a half media change at day 2.

### 2.4. Myelinating OL culture on aligned nanofibers

Passaged OPCs were seeded on 12 well aligned nanofiber inserts (Sigma Aldrich, #Z694614-12EA, fiber diameter = 700nm) at a density of 3*10^4^ cells/cm^2^ in OPC proliferation media, condition 3 of **Table 1a** The media was changed to OL differentiation media, condition 5 of **Table 1b** after two days of proliferation. Differentiation was maintained for four days with a half media exchange on the second day.

### 2.5. WST-8 Cell viability and proliferation assay

WST-8 assays were performed with OPCs cultured on 96 well plates, using the Quantimax^TM^ cell viability kit (BioMax, #QM1000) according to the manufacturer’s protocol. After isolation and proliferation for 72/96/120 hours, the media was aspirated, cells were washed with pre-warmed PBS, and 100_μ_L of DMEM without phenol red (Thermo Fisher Scientific, #21063029) was added. 10_μ_L of Quantimax^TM^ was promptly added to each well, and the plate was incubated for 30min in a 5% CO_2_ incubator. The optical density (OD) at 450nm was measured with a microplate reader. Proliferative capacity was calculated as relative values using the following formula: (450nm OD of well – average of 450nm OD values of blank)/(average of OD values of 450nm OD values of condition 1 of **Table 1a**.).

### 2.6. Immunofluorescence

Cells cultured on 9mm coverslips and nanofiber inserts were used for immunofluorescence. After designated culture periods, cells were fixed for 20 min with 4% paraformaldehyde and were stored at 4°C until use after washing three times with PBS. The cells were blocked with 10% normal goat serum (NGS), 0.1% Triton X-100 in PBS for 1 hour, RT. Cells were washed with PBS and were incubated overnight at 4°C with the following primary antibodies in the blocking solution; anti-NG2 (1:500, Millipore, #AB5320), anti-A2B5 (1:500, Millipore, #MAB312), anti-MBP (1:500, Abcam, #AB7349), anti-Olig2 (1:500, Merck Millipore, #MABN50), anti-GFAP (1:2000, Dako, #Z033429), anti-GFAP (1:2000, Abcam, #AB4674), anti-Tuj1 (1:1000, Dako, #G7121), anti-Iba1 (1:500, Wako, #019-19741), anti-cleaved-caspase-3 (1:200, Cell Signaling, #9664S), anti-Ki67 (1:500, Thermo Fisher Scientific, #MA514520). The cells were washed with PBS and incubated for 1hr, RT with appropriate Alexa Fluor 488-, 594-, 680-conjugated secondary antibodies (Thermo Fisher Scientific). Afterwards, the cells were counterstained with 4’,6-Diamidino-2-Phenylindole, dihydrochloride (DAPI, Sigma Aldrich, #D9542) for 10 minutes at RT and mounted onto glass slides. Cells were imaged with a Zeiss LSM800 confocal microscope.

### 2.7. EdU proliferation assay

EdU proliferation assays were performed with EdU Assay/EdU Staining Proliferation Kit (iFluor 488) (Abcam, ab219801) according to manufacturer’s protocol. Isolated OPCs were proliferated for 108 hours and incubated with EdU for 2 hours. Cells were fixed with 4% PFA and followed the immunofluorescence procedure, staining for Ki67. An additional EdU labeling step with the EdU reaction mix including iFluor 488 azide was applied between the secondary antibody incubation and DAPI counterstaining step, with an incubation period of 30 minutes at RT.

### 2.8. Statistics and image quantification

Independent t tests, one-way analysis of variance (ANOVA) with Tukey’s post hoc tests, simple linear regression, spline/LOWESS fitting, analysis of frequency distribution was performed with Graphpad Prism 9.5.1. *P*-values were two-sided, and a value of below .05 was considered as statistically significant. Cell counting was done using Fiji/ImageJ using the *Analyze particles* function when applicable. Measurements of nearest neighbor distances (NND) was made by first obtaining the XY coordinates of images of cell nuclei stained with DAPI in Fiji/ImageJ and converting the data table into a poisson point process using the spatial statistics (*spatstat*) package in R (version 4.2.0). Nearest neighbor distances and spatial G-function results were obtained using *spatstat*. Sholl analysis was performed using the *Neuroanatomy* plugin in Fiji/ImageJ. Myelin processes of OLs visualized with immunostaining for MBP, and binary masks were obtained by thresholding images. A center point at the location of the DAPI+ nuclei of the cell being analzyed was marked, and Sholl analysis was performed by drawing circles of increasing radius from the center point and quantifying the number of intersections between each circle and the binary mask. Comparison of morphological complexity was done by comparing the total number of intersections of cells. Schematics of culture procedure were created with BioRender.com.

## 3. Results

### 3.1. Isolation of OPCs from the neonatal rat brain

The development of the E3 method for primary OL culture started from an attempt to renovate a density gradient centrifugation-based method of isolating adult OPCs from the rodent cerebral cortex (Nihonmatsu-Kikuchi et al., 2021). Density gradient centrifugation has a major drawback, which is that the amount of tissue loadable into one column is substantially more limited compared to other centrifugation methods due to the concentration of cells and debris into one phase during the centrifugation process. To circumvent this problem, we adopted the principle of differential centrifugation, which exploits different sedimentation speed of particles and is widely used for cellular organelle fractionation or exosome isolation (Claude, 1946; Livshits et al., 2016) (**Fig. 1a**). In detail, after the dissection and dissociation of P1 rat cerebral cortices, the resulting cell suspension was passed through two sequential steps of centrifugation with the density gradient medium Optiprep^TM^ to exclude unwanted cells and debris. First, the cell suspension was centrifuged in a Hibernate-based dissection media with 12% Optiprep^TM^, and after centrifugation the supernatant was transferred to another 15mL tube, diluted to 6% Optiprep^TM^ (8mL in volume) and centrifuged again. The cell population from the pellet gained through the 2^nd^ centrifuge, when seeded at a low density of 1*10^4^cells/cm^2^ and cultured for 5 days in DMEM or DMEM/F12 based OPC proliferation media, exhibited colonies of NG2/A2B5+ OPCs comprising 90-95% of the total population, and a 3-5% proportion of background Tuj1+ neurons (**Fig. 1b, 1c, 1d and 1e**). GFAP+ astrocytes accounted for approximately 0.5% of the total population, and Iba1+ microglia were not observed **(Fig. S1b and S1c)**. In NBM-based proliferation media compositions, the OPC colonies showed reduced size and a lower proportion, approximately 75-77%. Additionally, Tuj1+ neurons accounted for approximately 20% of the total population. The cell pellet from the first centrifuge contained red blood cells, and when cultured with the same procedure showed a similar cell population, except for the fact that Iba1+ microglia could be discerned **(Fig. S1d)**. Thus, through two sequential density centrifugations, a cell population enriched in OPCs could be gained through 5-6 days of proliferation.

**Fig. 1.**
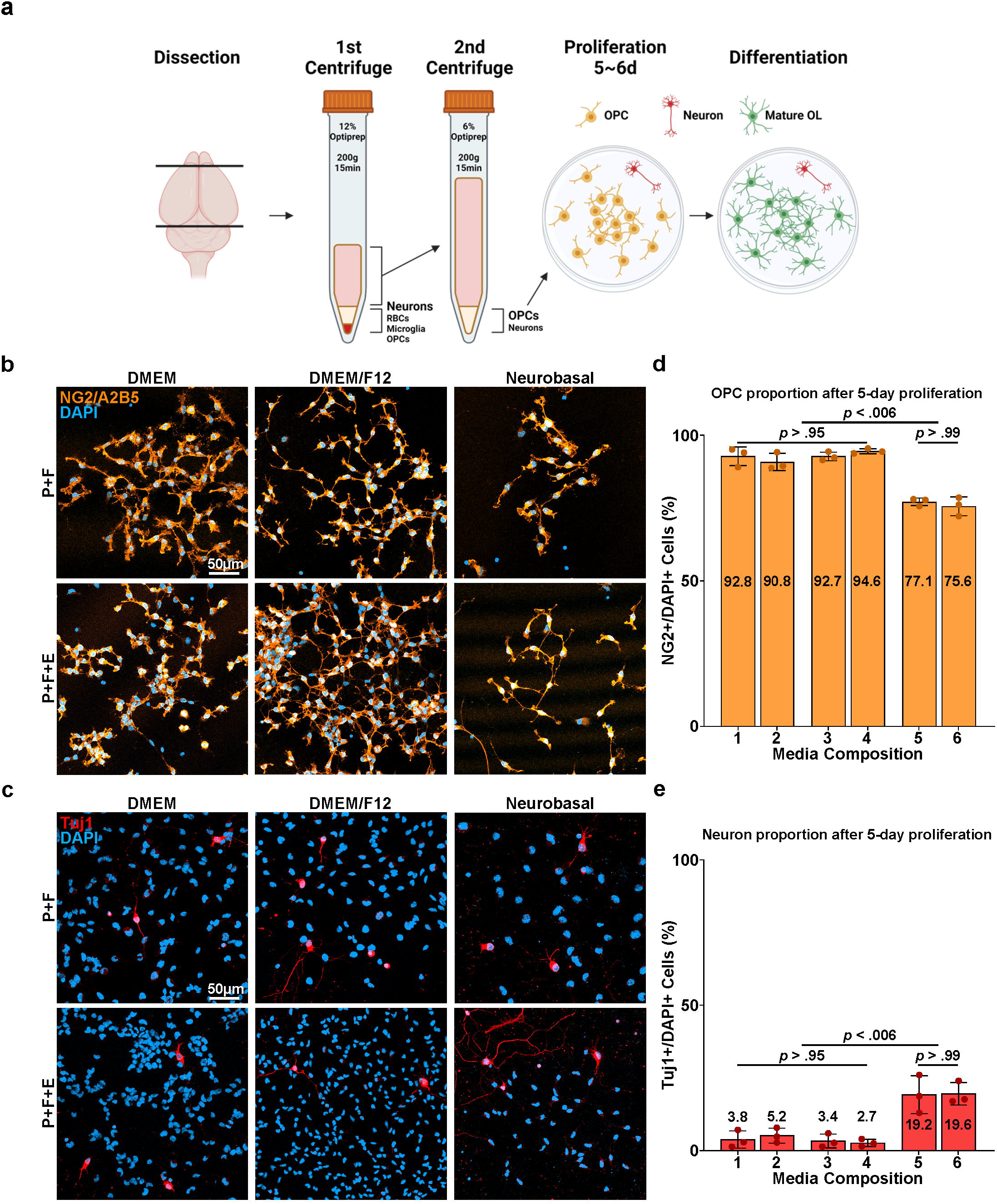
Cell composition of isolated OPC-enriched culture. **a** Scheme of the culture procedure. **b** Representative images of isolated cells (after 5 days of proliferation). Immunostaining for NG2 and A2B5. P: PDGF, F: FGF, E: EGF. **c** Representative images of isolated cells (after 5 days of proliferation). Immunostaining for Tuj1. P: PDGF, F: FGF, E: EGF. **d** Quantification of NG2+ OPC proportions after 5 days of proliferation according to media compositions corresponding to **Table 1a** (n=3, 5 images per media condition, approximately 300 DAPI+ cells per image. One-way ANOVA and Tukey’s post hoc test, two-sided. Data presented as mean values ± SEM.). **e** Quantification of Tuj1+ Neuron proportions after 5 days of proliferation according to media compositions corresponding to **Table 1a** (n=3, 5 images per media condition, approximately 300 DAPI+ cells per image. One-way ANOVA and Tukey’s post hoc test, two-sided. Data presented as mean values ± SEM.).

### 3.2. Media composition and growth factors for OPC proliferation

After the isolation of OPCs, various compositions of OPC proliferation media were compared to determine the most optimal formula to induce the proliferation of OPCs. The supplement B27, containing various nutritional factors and frequently used in serum-free oligodendrocyte culture media (O’Meara et al., 2011; Swire & Ffrench-Constant, 2019; Weil et al., 2019), was used in all conditions, as well as the growth factors PDGF and FGF, known to induce OPC proliferation and inhibit differentiation (Baron et al., 2000; Collarini et al., 1991; Hart et al., 1989; McKinnon et al., 1991; McKinnon et al., 1990). 6 conditions were compared based on differences in the base medium commonly used in primary OL culture: DMEM, DMEM/F12, NBM, and the presence/absence of EGF, which has been shown to promote OPC proliferation *in vitro* (Yang et al., 2017; Yang et al., 2016). Isolated OPCs were proliferated for 72∼120 hours in designated conditions. While immunostaining for the proliferation markers EdU and Ki67 showed no difference between media conditions in the proportion of cells proliferating, WST-8 proliferation/viability assays revealed that at both the 96- and 120-hour timepoint, condition 4 (DMEM/F12 with EGF) showed an approximately 10-15 fold higher viable cell number compared to NBM-based formulae. (**Fig. 2b, 2c and 2d**). Therefore, we selected condition 4, DMEM/F12 with EGF, as the optimal OPC proliferation medium, and following steps throughout the culture procedure were performed with OPCs proliferated in this media composition.

**Fig. 2.**
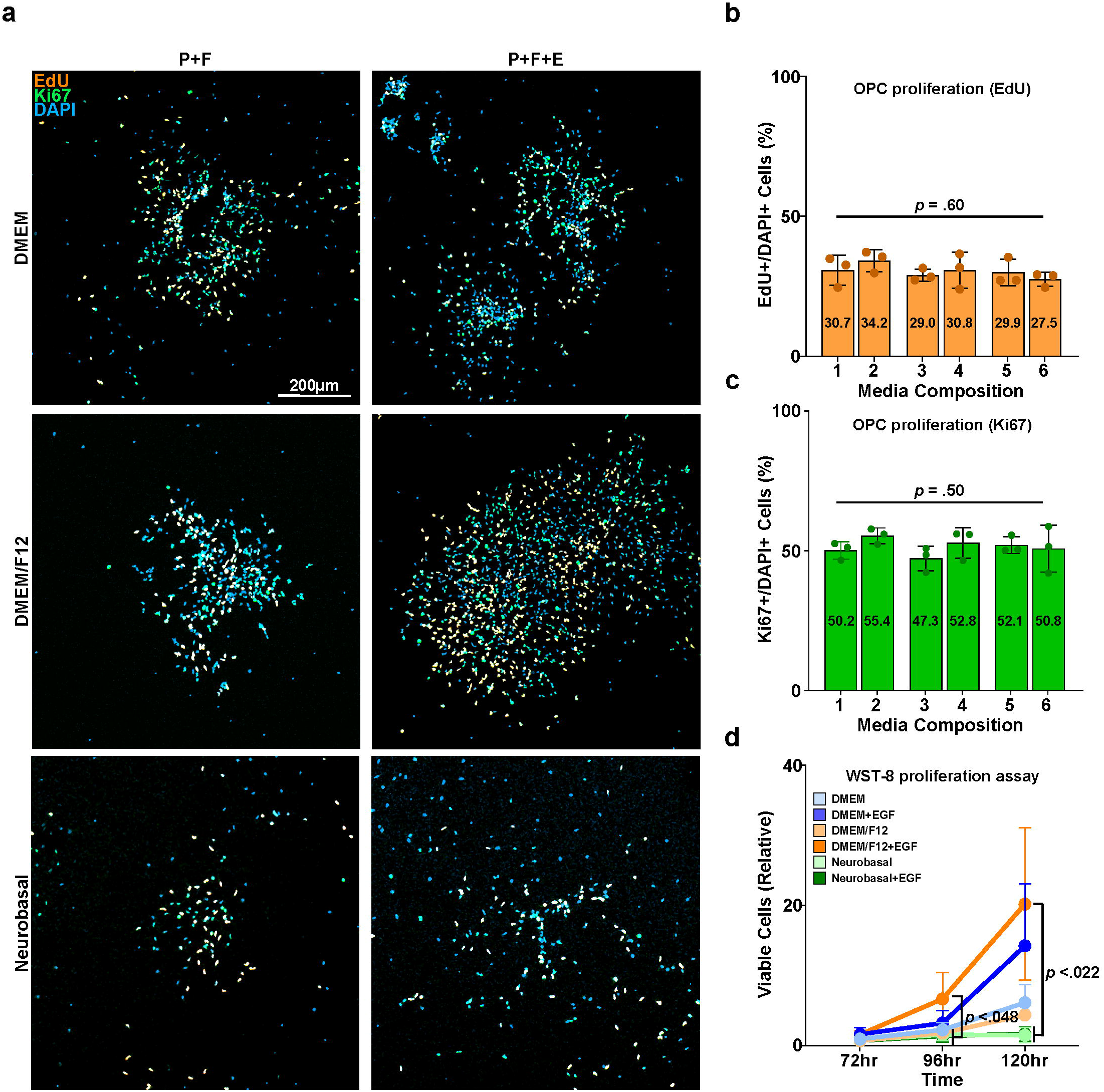
Proliferation of isolated primary OPC enriched culture. **a** Representative images of isolated cells (after 5 days of proliferation). Staining for EdU, Ki67. P: PDGF, F: FGF, E: EGF. **b** Quantification of EdU+/DAPI+ cells (%) according to media compositions corresponding to **Table 1a** (n=3, 10 images per media condition, approximately 500 DAPI+ cells per image. One-way ANOVA. Data presented as mean values ± SEM.). **c** Quantification of Ki67+/DAPI+ cells (%) according to media compositions corresponding to **Table 1a** (n=3, 10 images per media condition, approximately 500 DAPI+ cells per image. One-way ANOVA. Data presented as mean values ± SEM.). **d** Graph of WST-8 viability assay results, values relative to the DMEM (media composition 1) group cultured for 72 hours (n=3, 6 media conditions. One-way ANOVA and Tukey’s post hoc test, two-sided. Data presented as mean values ± SEM.).

### 3.3. Differentiation of OPCs into mature OLs and the passaging of proliferated OPCs

After successful proliferation of isolated OPCs, the culture media was exchanged to OL differentiation media. Different compositions of media, this time for the differentiation of OPCs into mature OLs, were tested. The use of B27 and N2 supplements which contain insulin like growth factor (IGF), known to be an essential factor for the survival of mature OLs *in vitro* (Barres et al., 1992), and thyroid hormone T3, a well-known OL differentiation-promoting factor, was uniform across all conditions (Ahlgren et al., 1997; Barres et al., 1994; Billon et al., 2002; Carre et al., 1998; Rodriguez-Pena, 1999). In a similar manner as OPC proliferation media conditions, another 6 conditions were tested, which could be distinguished by their base medium: DMEM, DMEM/F12, NBM, and the presence/absence of CNTF, a growth factor known to promote OL maturation and thus commonly included in OL differentiation media formula (Stankoff et al., 2002; Talbott et al., 2007). After 4 days of differentiation, MBP+ mature oligodendrocytes with web-like myelin sheets could be observed in all groups (**Fig. 3b**). However, aside from the successful differentiation of OPCs into mature OLs, we discovered areas of clumped oligodendrocytes with entangled MBP+process indistinguishable from one another, showing nuclear condensation and immunoreactivity for cleaved caspase 3, indicating they had undergone apoptosis (**Fig. 3c**). When visualized at low magnification, such clumps of apoptotic OLs were most prominent in the center of colonies where focal cell density is highest (**Fig. 3a**). To confirm the association between high cell density and cell death during differentiation, we analyzed the correlation between the average nearest neighbor distance of cells and the rate of cleaved caspase-3 positive dead cells in images containing 100-600 cells per image. We discovered an inverse correlation between mean nearest neighbor distance and cell death rate, showing that the overcrowding of OPCs during proliferation caused cells to die during induction of maturation (**Fig. 3d**). An additional procedure to redistribute proliferated OPCs and acquire more space was deemed necessary.

**Fig. 3.**
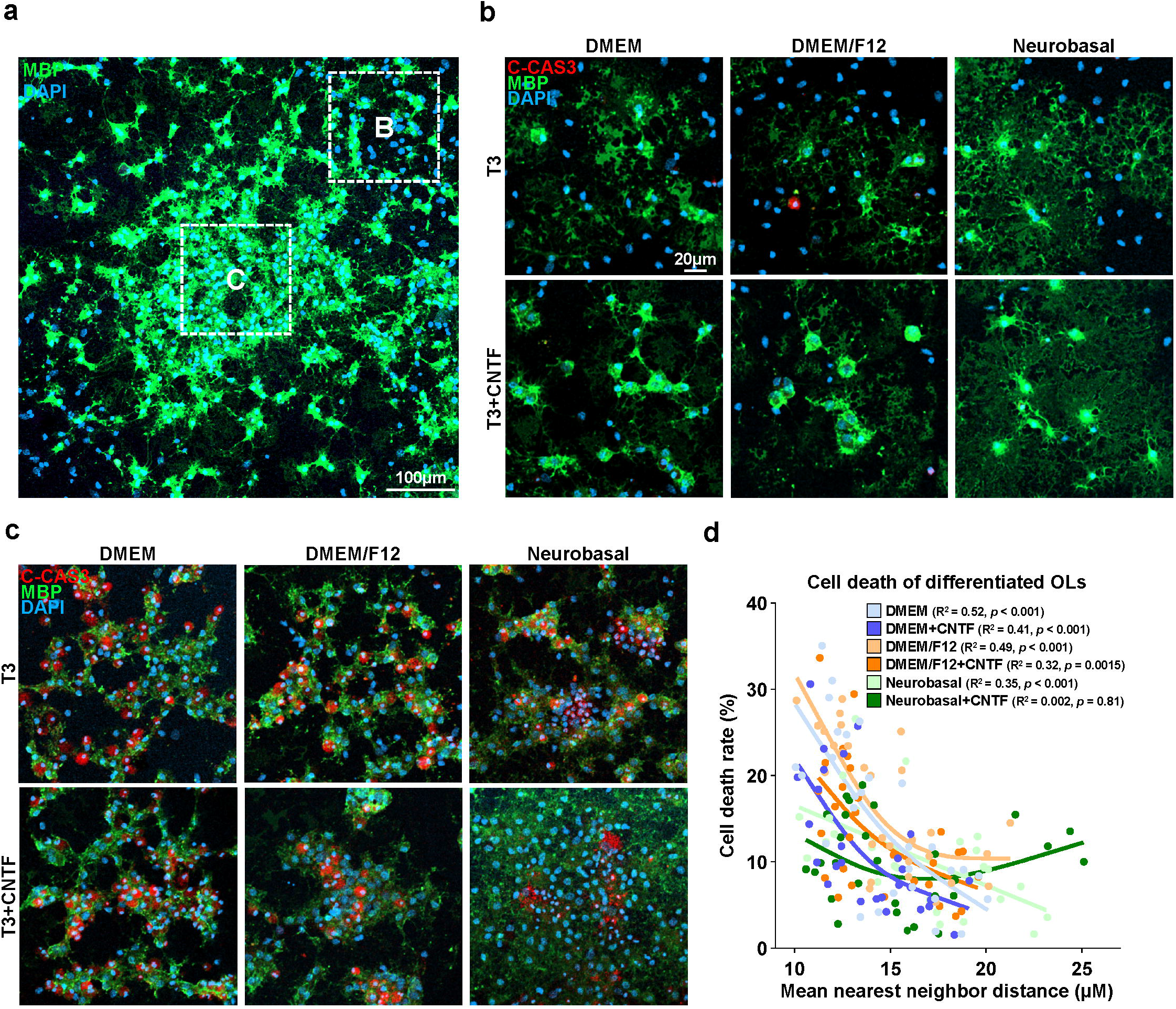
Differentiation of isolated OPC-enriched culture. **a** Representative images of isolated cells after 4 days of differentiation. Immunostaining for myelin basic protein (MBP). **b** Representative images of isolated cells after 4 days of differentiation, in the periphery of colonies (Area B of **Fig. 3a**). Immunostaining for cleaved-caspase 3, MBP. **c** Representative images of isolated cells after 4 days of differentiation, in the center of colonies (Area C of **Fig. 3a**). Immunostaining for cleaved-caspase 3, MBP. **d** Scatter plot of the mean nearest neighbor distance of cells in immunostained images and cell death rate measured as cleaved caspase-3+/DAPI+ cells (%), and spline/LOWESS fitting. The correlation between mean nearest neighbor distance and cell death rate was analyzed with simple linear regression for each media condition (n=3, 10 images per media condition, approximately 300 DAPI+ cells per image).

For the redistribution and spacing of OPCs to avoid cell death during differentiation, proliferated OPCs were passaged and redistributed, seeded at a density of 1*10^4^cells/cm^2^ (**Fig. 4a**). Passaged OPCs were stabilized for 2 days in OPC proliferation media. The previous 6 proliferation media formulas were tested again to confirm which formula was most adequate for the 2-day stabilization period. We observed that after passaging, the background Tuj1+ neurons which occupied 5% of the total population during OPC proliferation was now reduced to a near-nonexistent state, and the purity of Olig2-positive OL lineage cells, OPCs judging from their bi/tripolar morphology and immunoreactivity for NG2, was increased from the prior 90-95% to approximately 98-99% (**Fig. S2a, S2b, S2b and S2c)**. The average nearest neighbor distance of passaged cells after 4 days of differentiation was increased from approximately 15_μ_m to 19_μ_m compared with cells differentiated without passaging (*P* < .001, nested independent t-test, n=3, two-sided), and the mean cell death rate after 4 days of differentiation, measured as the percentage of cleaved caspase-3 positive cells was reduced from 13% to 4% (*P* < .001, nested independent t-test, n=3, two-sided) (**Fig. 4f and 4g**). We observed that use of DMEM/F12 with EGF, which was the formula of choice during the OPC proliferation period before redistribution, appeared to cause cells to have a more inhomogeneous spatial distribution compared to other media conditions. To quantify the “randomness” of spatial distribution, The coordinates of cell nuclei shown by DAPI in a square image was converted to Poisson point processes, and histograms of nearest neighbor distances and the G-function, a cumulative distribution function of nearest neighbor distances were drawn. While cells stabilized after passage in the other 5 media compositions showed a distribution within the boundaries of complete spatial randomness, cells stabilized in DMEM/F12 with EGF showed a left shift of both the histogram of NND frequency distribution and the G function, indicating clustering of cells (**Fig. 4d and 4e**). Cells showing a morphology differing from typical OPCs were also visible in media conditions including EGF, especially when combined with NBM, and therefore DMEM/F12 without EGF, condition 3 of **Table 1a** was selected as the media for the post-passage stabilization period (**Fig. 4b**).

**Fig. 4.**
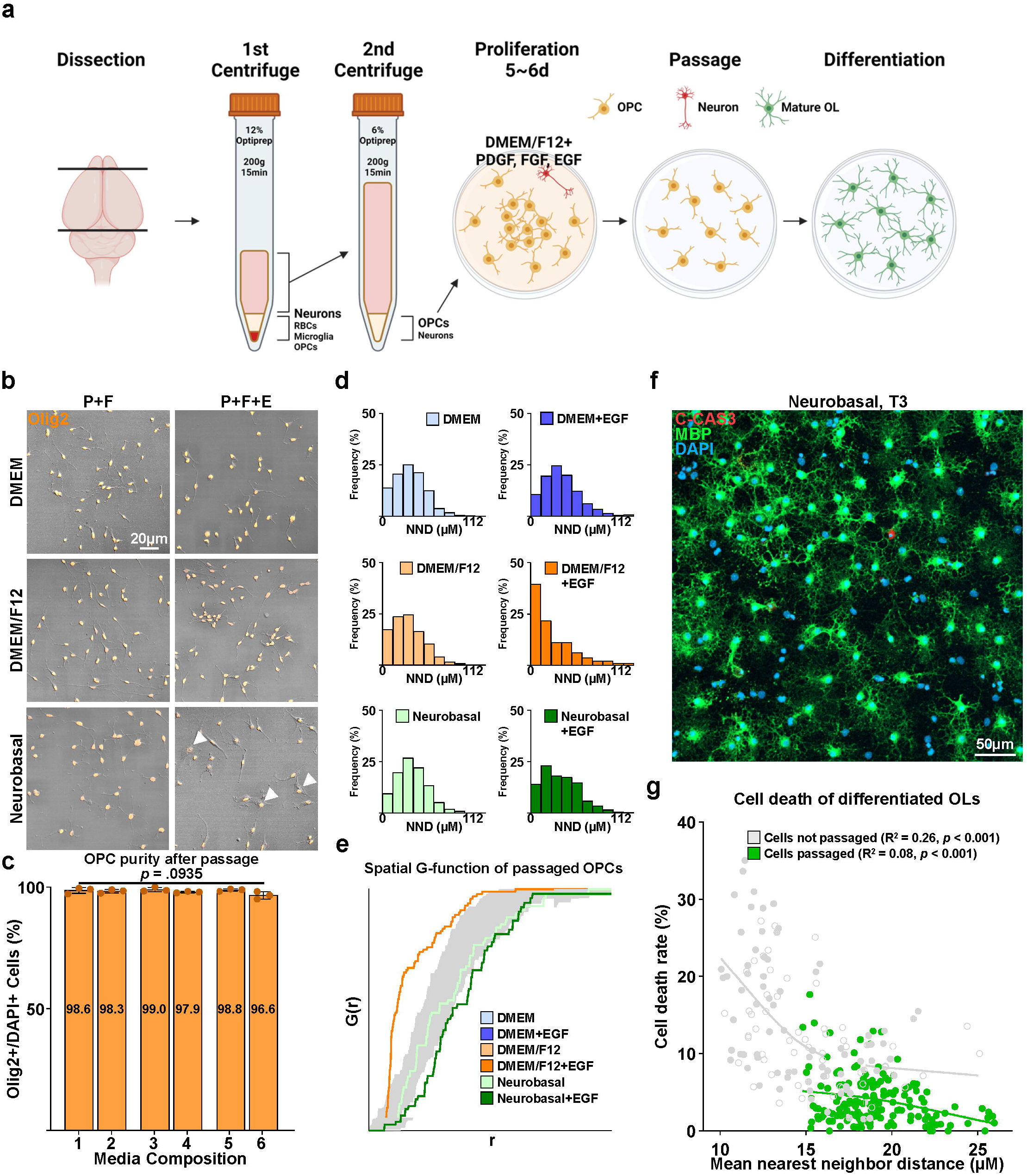
Characterization of passaged OPC culture. **a** Scheme of revised culture procedure, involving the passaging of expanded OPCs. **b** Representative images of OPCs passaged and redistributed. Immunostaining for Olig2. Arrowheads; Olig2+ cells with atypical morphology. **c** Quantification of Olig2+/DAPI+ cells (%) according to media compositions corresponding to **Table 1a** (n=3, 10 images per media condition, approximately 50 DAPI+ cells per image. One-way ANOVA. Data presented as mean values ± SEM.). **d** Representative histograms of nearest neighbor distances (NND) of OPCs after passaging (n=3, 16 images per media condition, approximately 200 DAPI+ cells per image). **e** Graph of spatial G-function (cumulative distribution of nearest neighbor distances) of passaged OPCs by proliferation media composition. Gray zones demarcate boundaries of complete spatial randomness. **f** Representative low-magnification image of passaged OPCs after 4 days of differentiation into mature OLs in a Neurobasal-based differentiation media (Condition 5 of **Table 1b**). Immunostaining for cleaved caspase 3, MBP. **g** Scatter plot of the mean nearest neighbor distance of cells in immunostained images and cell death rate measured as cleaved-caspase 3+/DAPI+ cells (%), and spline/LOWESS fitting. The correlation between mean nearest neighbor distance and cell death rate was analyzed with simple linear regression for each condition (n=3, 9 images per media condition.), approximately 200 DAPI+ cells per image).

### 3.4. Differentiation of passaged OPCs into mature OLs

After passaged OPCs had been stabilized and proliferated for 2 days, we once again tested the six OL differentiation media formulas. After 2 days of differentiation, MBP positive mature OLs with branch-like processes could be observed, comprising around 35∼50% of the population regardless of the media used (**Fig. 5a and 5b**). By the 4^th^ day, web-like myelin sheets could be observed in all groups. Despite the difference in levels of maturation, the actual number of MBP+ OLs were not significantly different between cultures differentiated for 2 and 4 days (**Fig. 5a and 5b**). To compare the degree of differentiation by morphological complexity, Sholl analysis was performed. Results revealed that NBM-based formulas produced OLs with significantly more complex processes at day 4 (**Fig. 5c**). The use of CNTF did not influence OL maturation in terms of number or morphological complexity (**Fig. 5a, 5b and 5c**). Overall, we successfully induced the differentiation of OPCs into mature OLs and discovered that NBM-based differentiation media greatly enhanced the morphological complexity of mature OLs.

**Fig. 5.**
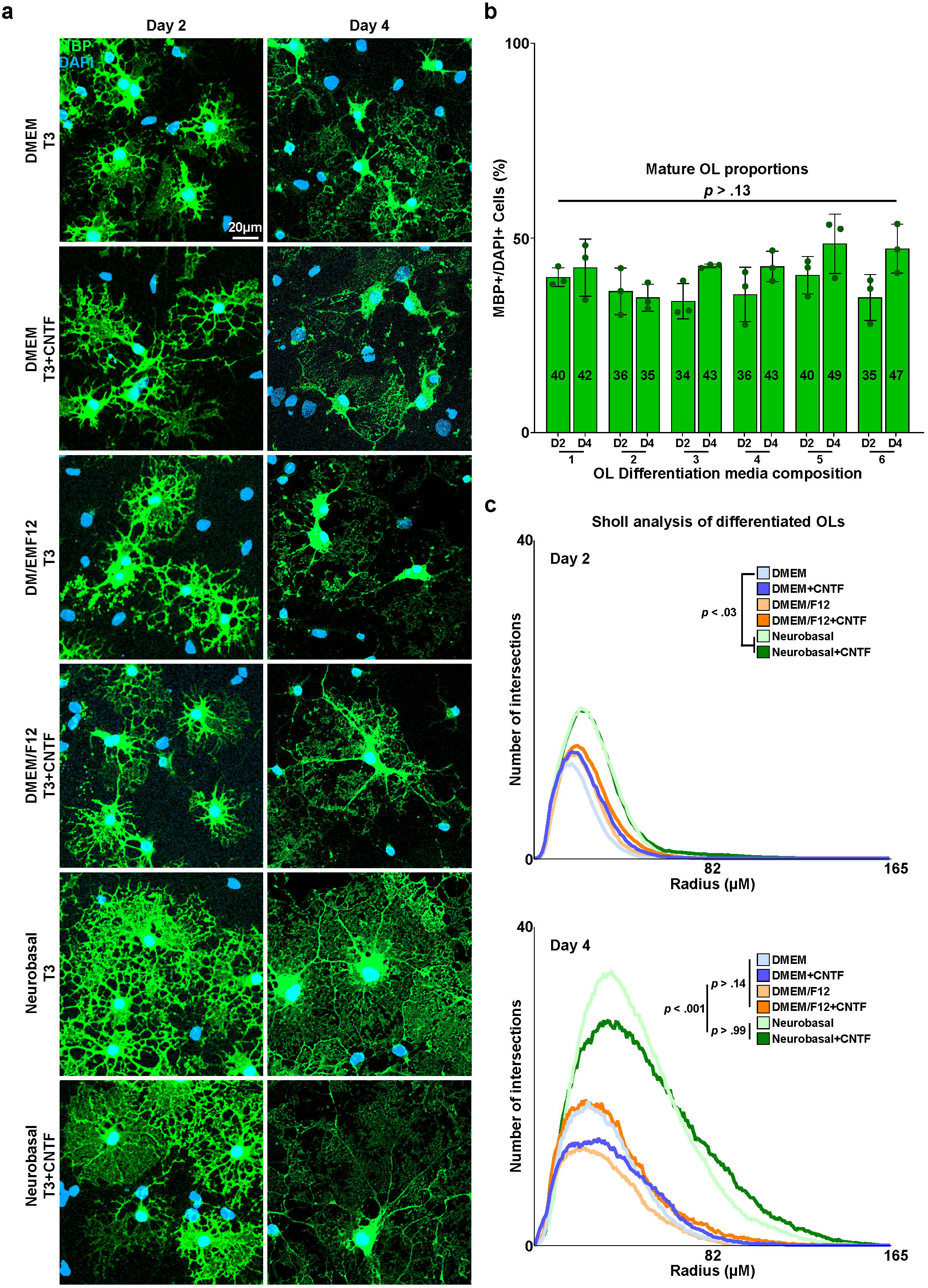
Differentiation of passaged OPC culture. **a** Representative images of differentiated OLs, differentiated after passaging for 2 or 4 days in designated OL differentiation media compositions. Immunostaining for MBP. **b** Quantification of MBP+/DAPI+ cells (%) of passaged OPCs after designated differentiation periods and media compositions corresponding to **Table 1b**. (n=3, 8-10 images per media condition, approximately 60 DAPI+ cells per image. One way ANOVA and Tukey’s post hoc test, two-sided. Data presented as mean values ± SEM.). **c** Results of Sholl analysis, of differentiated OLs. Comparison of morphological complexity was performed by comparing the total number of intersections per cell. (n=3, 10 images per media condition, approximately 10 MBP+ OLs per image. One way ANOVA and Tukey’s post hoc test, two-sided. Data presented as mean values ± SEM.)

### 3.5. Myelin sheath formation of OLs culture through the E3 method

Lastly, after confirmation of successful OL differentiation, we validated the capacity of cultured and maturated OLs produced with the E3 method to enwrap neurons or neuron-like structures and form myelin sheaths (Bechler, 2019; O’Meara et al., 2011; Swire & Ffrench-Constant, 2019; Yoshida et al., 2020). Having observed the presence of background neurons during OPC proliferation after isolation, primary neuron cultures were obtained by seeding isolated cells from the 2^nd^ centrifuge step with 6% Optiprep^TM^ at a high density of 5*10^4^cells/cm^2^ in NBM supplemented with B27 and N2, without the OPC favoring growth factors PDGF and FGF. After culture for 14 days, robust axonal processes could be observed. We seeded passaged OPCs on the neuron cultures at a density of 3*10^4^cells/cm^2^, and the media was exchanged to differentiation media condition 5, NBM-based media without CNTF. At day 4 of differentiation, we discovered that areas of overlapping Tuj1 positive axons and MBP positive myelin sheaths were indeed present (**Fig. 6a**). Furthermore, we also examined the ability of cultured OLs to enwrap aligned nanofibers. The whole culture procedure from isolation to differentiation was repeated, except that passaged OPCs were seeded on 12-well aligned nanofiber inserts of 700nm diameter. Z-stack images of MBP positive myelin sheaths on the aligned fibers visualized by the differential interference contrast (DIC) filter were taken, and we noted efficient formation of myelin sheaths in 3D reconstructions, which were wrapped around the aligned fibers (**Fig. 6b**). Thus, we confirmed that OLs cultured through the E3 method could form myelin sheaths and enwrap neurons *in vitro*.

**Fig. 6.**
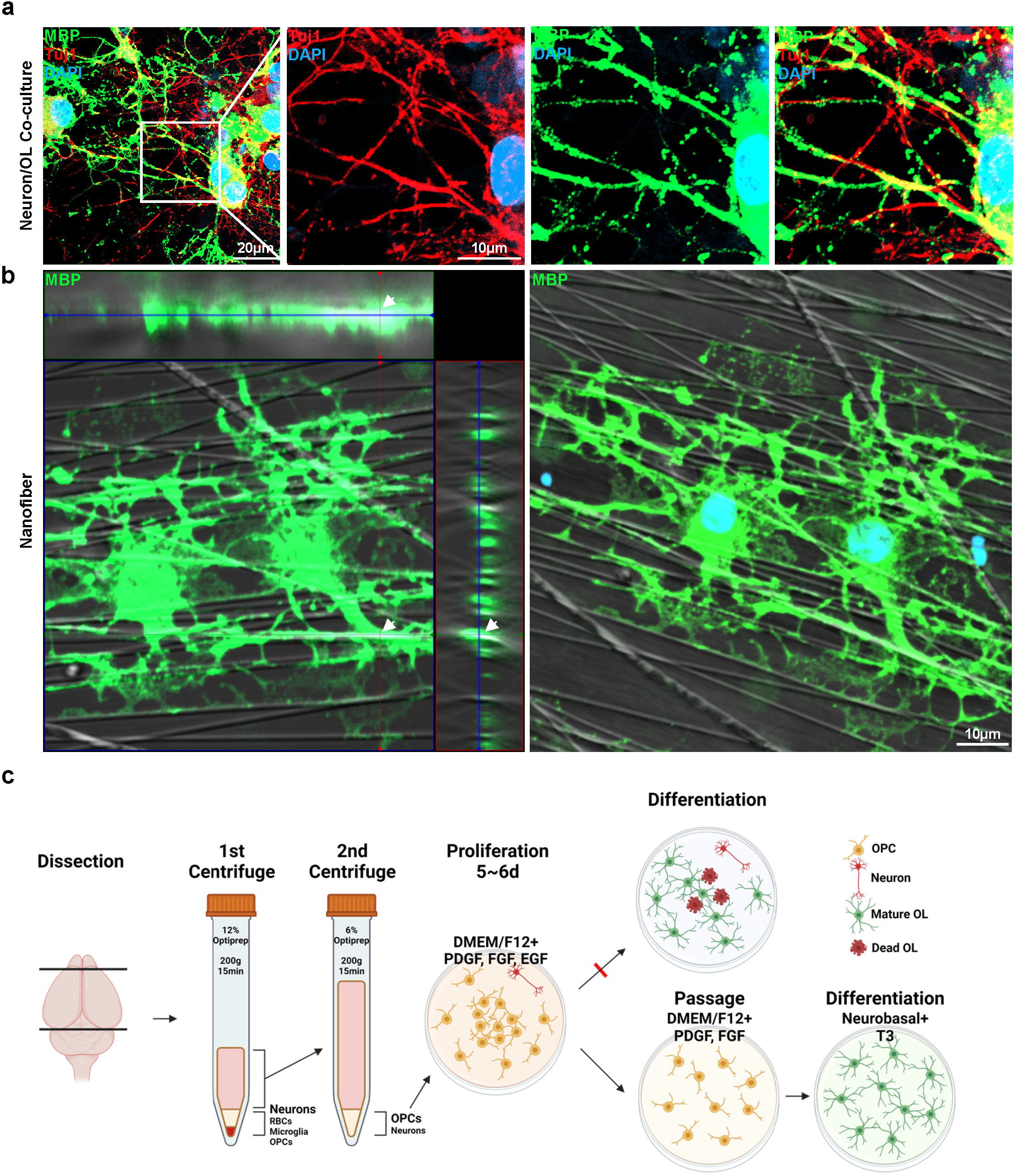
Final culture procedure and myelination capacity of cultured oligodendrocytes. **a** Representative images of neuron/oligodendrocyte co-culture. Immunostaining for MBP, Tuj1. **b** Representative images of differentiated oligodendrocytes cultured on aligned nanofibers. Immunostaining for MBP**. c** Final schematic of the full culture procedure.

## 4. Discussion

Research on oligodendrocytes and myelination greatly benefit from *in vitro* culture models, due to the deconvolution of numerous and dynamic stages of the OL lineage. All stages of the OL lineage, even OPCs, are present in the brain throughout life, and the stage of a typical OL lineage cell is not constant, being subject to change. This may make even simple observations such as the counting of cell number complicated, and dependent on the use of multiple stage-specific reporters. Such considerations emphasize the need for an efficient method for *in vitro* primary OL culture. We introduce the E3 method for primary OL culture through the present study, which greatly improves the efficiency of procedure and cell yield.

The shaking method, first developed by McCarthy, is the oldest and most widely used method for isolating primary OPCs (Choi et al., 2017; Choi et al., 2023; McCarthy & de Vellis, 1980; O’Meara et al., 2011; Yang et al., 2005; Zhu et al., 2014). This method exploits the different adhesion capacity of glial cell types to shake off OPCs from a mixed glial culture including animal serum. Despite the advantage of a simple procedure, the shaking method has a few drawbacks: The use of serum, a long mixed glial culture period of 7∼10 days, and potential mechanical injury to cells due to overnight shaking (**Table 2**.). Antibody based methods, especially immunopanning, has the advantage of the direct acquisition of OPCs from tissue without a separate expansion process and a high purity above 99.5%, but a prolonged dissection time due to multiple panning steps and expensive requirements such as antibodies for the negative and positive selection of cells, and hybridomas for the generation of antibodies are required to achieve results (Barres et al., 1992; Dugas & Emery, 2013a, 2013b; Emery & Dugas, 2013; Mayer-Proschel, 2001) (**Table 2**.). More recent methods based on density gradient centrifugation partially address problems found in conventional methods, but have the disadvantage of a low cell yield due to the “clogging” of columns caused by concentration of cells and debris into one plane during centrifugation, which places a limit on the amount of tissue processable per column (Brewer & Torricelli, 2007; Nihonmatsu-Kikuchi et al., 2021). OPC isolation in the E3 method is based on differential centrifugation, requiring only the iodixanol density gradient medium Optiprep^TM^, and the augmentation of OPC colonies has been achieved with a serum-free proliferation medium comprising the three growth factors PDGF, FGF and EGF. We shortened the period of OPC growth to 5 days, and minimized damage to cells from an overnight shaking procedure to a 5∼10-minute detachment with Accutase®, which also improved the OPC purity to a number comparable to the shaking method (Choi et al., 2017; Choi et al., 2023; McCarthy & de Vellis, 1980; O’Meara et al., 2011; Yang et al., 2005; Zhu et al., 2014) (**Table 2**). The quality of the cultures was comparable at the OPC (shaken vs passaged) and differentiated mature OL stage **(Fig. S3).**

**Table 2.** Comparison of the E3 method with conventional primary OL culture methods.

| Method<br>Parameter | Shaking | Immunopanning | E3 |
| --- | --- | --- | --- |
| **Dissection time**<br>( Sacrifice ~ incubation ) | **Approx. 2 hours**<br>3min / pup dissection<br>30-60min dissociation<br>+ $\alpha$ | **Approx. 4 hours**<br>3min / pup dissection<br>30-60min dissociation<br>15-45min panning*3<br>+ $\alpha$ | **Approx. 2 hours**<br>3min / pup dissection<br>30-60min dissociation<br>15min centrifuge*2<br>+ $\alpha$ |
| **Time required for acquisition of OPCs** | **7-10 days**<br>Time until shaking | **Approx. 4 hours**<br>Dissection time | **5-6 days**<br>Time until passage |
| **Purity of OPCs** | **90~98%**<br>Contaminant =<br>Astrocytes, Microglia | **>99.5%**<br>Contaminant =<br>Fibroblast-like cells | **98-99%**<br>Contaminant =<br>Neurons |
| **Exclusive materials required** | **Fetal Bovine Serum**<br>**Shaking Incubator** | **Antibodies**<br>( anti-RAN2, anti-BSL1,<br>anti-GalC, anti-O4,<br>anti-A2B5, anti-PDGFRα )<br>**Hybridomas** | **Iodixanol (Optiprep™)** |
| **References** | (McCarthy & de Vellis, 1980)<br>(Yang et al., 2005)<br>(O'Meara et al., 2011)<br>(Zhu et al., 2014)<br>(Choi et al., 2017)<br>(Choi et al., 2023) | (Barres et al., 1992)<br>(Mayer-Proschel, 2001)<br>(Dugas & Emery, 2013a)<br>(Dugas & Emery, 2013b)<br>(Emery & Dugas, 2013) |  |

During the development of the E3 method, we discovered a few intriguing *in vitro* characteristics of OL lineage cells, which may provide implications for both *in vitro* and *in vivo* research on OL physiology and pathology. The first is the dual effect of NBM on OPC proliferation and OL differentiation. We found that NBM showed a significantly worse performance compared to DMEM or DMEM/F12 in terms of OPC proliferation, but when it came to the differentiation of OPCs into mature OLs, they produced OLs with much more ramified myelin processes. Being a media originally developed for the culture of primary neurons, the most prominent feature which separates NBM from both DMEM and DMEM/F12 is its low osmolarity of around 235mOsm when combined with the B27 supplement, compared to the osmolarity DMEM or DMEM/F12, which is around 335mOsm (Brewer et al., 1993). There exists a previous study which shows that heightening the osmolarity of NBM with NaCl or mannitol boosts its capacity for OPC proliferation to levels comparable with DMEM (Kleinsimlinghaus et al., 2013). Prior research explicating the effect of extracellular matrix (ECM) stiffness and mechanosensing on the proliferation and differentiation of OPCs are also present (Domingues et al., 2018; Segel et al., 2019), which may be a possible mechanism behind the effect of the low osmolarity of NBM, taking into consideration that cells can sense osmolarity through detection of mechanical cell membrane stretching due to different osmolarity levels. The results of our study suggest that the low osmolarity of NBM also contributes to a better induction of OL maturation. The dual effect of osmolarity on OL dynamics may not be limited to *in vitro* systems and may possibly play a role *in vivo* such as during developmental myelination or remyelination after injury, and further research on the influence of osmolarity levels on OPC proliferation and OL maturation is necessary.

Another notable feature of OLs we discovered was the death of cells during differentiation caused by the overly close distribution of OPCs. OPCs in the center of colonies formed during proliferation developed into entangled OLs impossible to discern from each other, and a large portion of these cells were positive for apoptosis markers. An inverse correlation between the average space between OPCs and cell death during induction of differentiation could be seen. The formation of overcrowded OPC colonies hint at their stemness, which is more obvious in the formation of “oligospheres” from neural progenitors observed *in vitro (Avellana-Adalid et al., 1996; Vitry et al., 1999; Zhang et al., 1998)*. Although the degree of OPC clustering shown in the E3 method or oligospheres may be a phenomenon limited to *in vitro* circumstances, there are numerous past studies on the reactive proliferation of OPCs after injury and their insufficient differentiation and remyelination (Back, 2017; Back et al., 2002; Susarla et al., 2014). We speculate that the overcrowding of OPCs may cause insufficient differentiation and cell death. Also, the association of OPC clustering and cell death may provide points to contemplate for research on OPC transplantation to promote remyelination after injury. Numerous studies on OPC transplantation have been conducted, and they have been shown to have a meaningful effect (Li et al., 2021; Sun et al., 2013; Wang et al., 2020; Wang et al., 2021; Wu et al., 2012; Xu et al., 2015). However, in many cases an excessively large number of OPCs are grafted to the site, which may recreate the situation of OPC crowding discerned in our study. The findings ascertained during the development of the E3 method may provide information on the appropriate density of OPCs for successful transplantation.

We also tested the use of growth factors commonly used in primary OL culture. Growth factors more or less unanimously used, such as PDGF and FGF for OPC proliferation, and IGF and T3 for OL differentiation were fixed, and we evaluated the effect of factors less known and used compared to the factors listed above, which were EGF for the OPC stage and CNTF for the mature OL stage. The use of EGF during the proliferation of OPCs first isolated from the rat brain enhanced the proliferation of OPCs, outperforming media compositions without EGF in WST-8 proliferation assays. Also, when applied to passaged OPCs for 2 days, it caused cells initially seeded in a random pattern to rearrange into a clustered appearance. Taking into accord the fact that the cell number per area were not different in conditions with EGF compared to those without (data not shown), the most plausible explanation for this clustered arrangement is that OPCs were more migratory in an EGF-included environment. Past studies on EGF and oligodendrogenesis propose that EGF promotes both the proliferation and differentiation of OPCs, and there is a previous study which suggests that EGF causes OPCs to transit into glial progenitor cells (GPC) (Gonzalez-Perez & Alvarez-Buylla, 2011; Yang et al., 2017; Yang et al., 2016). Although our study does not provide evidence on the presence of GPCs, the proliferation and migration-promoting property of EGF in our proliferation media compositions does strongly imply that EGF keeps OPCs in a more stem cell-like state. As for the use of CNTF, we did not observe a significant enhancement in OL maturation.

In conclusion, we introduce the E3 method of primary OL culture, optimizing existing methods for OPC isolation and media compositions for subsequent proliferation and differentiation. This method retains the simplicity of procedure, while producing improved results in aspects of both efficiency and quality, compared to widely used methods of OL culture. Also, *in vitro* features of the OL lineage discovered during development of the E3 method may provide suggestions for research on oligodendrocyte physiology and pathophysiology.

## Funding information

This research was supported by a grant of the M.D.-Ph.D./Medical Scientist Training Program through the Korea Health Industry Development Institute (KHIDI), funded by the Ministry of Health & Welfare, Republic of Korea (to H. K.). This work was also supported by National Research Foundation of Korea (NRF) grants funded by the Korean government (MSIT; Ministry of Science and ICT) (NRF2019R1A5A2026045 and NRF-2021R1F1A1061819), a grant from the Korean Health Technology R&D Project through the Korea Health Industry Development Institute (KHIDI), funded by the Ministry of Health & Welfare, Republic of Korea (HR21C1003), and new faculty research fund of Ajou University School of Medicine (to J.Y.C.).

## Data availability statement

The data and analyses in this study are available within the main text and the Supporting Information section. The R code used for spatial analysis of poisson point processes are available under https://github.com/hanki313/Spatial-statistics-for-E3-methods/tree/main

## Supporting information

Protocol of culture method

## Acknowledgments

Fig. 1a, 4a and 6c were created with Biorender.com. We thank Jonah R Chan from the University of California, San Francisco, for advice on designing the *in vitro* nanofiber myelination experiments.

## Notes

### Competing Interest Statement

The authors have declared no competing interest.

