## Supplementary material for "A primary culture method for the easy, efficient and effective acquisition of oligodendrocyte lineage cells": Protocol of culture method

**Protocol for E3 primary oligodendrocyte culture**

**Preparements**

**Reagents**

- Postnatal day 1 Sprague-Dawley rat pups
- Poly-D-Lysine (PDL, Sigma Aldrich, #P6407) solution
- 70% Ethanol (in distilled water)
- Hank’s Balanced Salt Solution (HBSS, no Calcium, no Magnesium)
- Phosphate Buffered Saline (PBS, no Calcium, no Magnesium)
- Accumax^TM^ (Merck Millipore, #SCR006) dissociation enzyme
- Accutase® (Thermo Fisher Scientific, #A11105-01)
- Papain suspension (Worthington, #LS003126)
- Optiprep^TM^ (Sigma Aldrich, #D1556) density gradient medium
- Hibernate-A medium (Thermo Fisher Scientific, A1247501)
- Dulbecco’s modified eagle medium (DMEM, Cytiva, #SH3002201)
- Dulbecco’s modified eagle medium/Ham’s F12 (DMEM/F12, Thermo Fisher Scientific, #11320033)
- Neurobasal medium (NBM, Thermo Fisher Scientific, #21103049)
- Penicillin/streptomycin (Cytiva, #SV30010)
- GlutaMAX^TM^ (Thermo Fisher Scientific, #35050-061)
- B27 supplement (Thermo Fisher Scientific, #17504-044)
- N2 supplement (Thermo Fisher Scientific, #17502048)
- Platelet derived growth factor-AA (PDGF-AA, Peprotech #100-13A)
- Basic fibroblast growth factor (bFGF, Peprotech #100-18B)
- Epidermal growth factor (EGF, Peprotech #AF-100-15)
- Triiodothyronine (T3, Sigma, #T6397)

**Equipment**

- T75 cm^2^ culture flasks with vent-seal (Corning)
- Culture surfaces (6- ,12- well plates, Corning)
- Conical tubes (50mL, 15mL, SPL)
- 10cm Petri dishes (SPL)
- Dissection Instruments: Surgical scissors (Large and small), curved forceps (#7), forceps
- Laminar flow hood
- Humidified culture incubator (37℃, 5% CO_2_)
- Water bath at 37℃, 30℃
- Tabletop centrifuge
- Equipment for cell counting
- Bright field microscope

**Solutions and media**

- Dissection medium: Hibernate-A medium, 1% GlutaMAX^TM^, 1% Antibiotics
- Dissociation medium: 4mL Dissection medium, 100uL papain suspension (per animal = brain)
- 12% Optiprep^TM^ medium: 4mL Dissection medium, 12% Optiprep^TM^ (per animal = brain)
- Oligodendrocyte progenitor cell (OPC) medium: DMEM/F12, 1% GlutaMAX, 1% Penicillin/streptomycin, 2% B-27 supplement, 30ng/mL PDGF-AA, 10ng/mL FGF, 10ng/mL EGF
- Oligodendrocyte (OL) differentiation medium: Neurobasal media, 1% GlutaMAX, 1% Antibiotics, 1% N2 supplement, 2% B-27, 40ng/mL T3

**Procedure** (If not specified otherwise, for one rat pup (brain))

***Preparation for culture***

1. Coat cell culture surfaces in 10µg/mL PDL (for the quoted product, 1:10 dissolved in distilled water). A minimum of 1 hour coating is necessary, and surfaces must be dried before use.
2. Incubate dissociation media in a 30°C water bath for 30 minutes to activate the enzyme.

*Although the exact optimal temperature for the activation of papain is known to be 30°C, 37°C can be used for convenience.

***Dissection of P1 rat brain***

3. Place the rat pup(s) on ice for anesthetization, and after the pups have been anesthetized, decapitate with surgical scissors (large), and place the heads in a 50mL conical tube containing ice-cold HBSS.
4. Transfer the head(s) to another 50mL conical tube with ice-cold 70% Ethanol, gently tap the tube, and transfer the heads to another tube containing fresh ice-cold HBSS.
5. Transfer a head to a petri dish with ice-cold HBSS.
6. Using the blunt side of the curved forceps, grasp (pinch) the scalp at the midline of the cranium and tear it off.
7. Cut the now-exposed skull with surgical scissors (small) along the midline superior sagittal sinus and tear them off with curved forceps.
8. Severe the connection between the olfactory bulb and the rest of the brain.
9. Advance the curved forceps beneath the cerebellum and scoop out the brain.
10. Isolate the two cerebral hemispheres, remove the meninges (A nick in the meninges can usually be seen by the gross eye or found near the rostral portion of the hemisphere where the olfactory bulb was severed off.), and place the hemispheres in a 15mL conical tube containing 1mL of ice-cold Accumax^TM^. Repeat steps 5–10 until all pups have been processed.

*Hence, all steps should be performed within a laminar flow hood when possible.

***Dissociation of tissue with Papain***

11. Briefly triturate the brain(s) 10 times with a 1mL pipette tip.
12. Add 4mL of the dissociation solution and incubate for 30 minutes at 30°C or 37°C.

***Differential centrifugation with Optiprep^TM^***

13. Centrifuge the cell suspension(s) at 200g for 5minutes. Aspirate the supernatant and resuspend in 1mL of 12% Optiprep^TM^ medium.

*Inactivation of papain can be performed but isn’t necessary. We advise against using serum for inactivation, as serum can affect the fate commitment of OPCs.

14. Triturate 10-20 times with a 10µL pipette tip attached to a 1mL tip. Add 3mLs of 12% Optiprep^TM^ medium to obtain a 4mL cell suspension in 12% Optiprep^TM^ medium.

*All trituration steps should be carefully performed to prevent introducing bubbles in the cell suspension(s)

15. Centrifuge the cell suspension(s) at 200g, 15min at room temperature (RT), full brake.
16. Transfer the supernatant (4mL) to a different 15mL tube and add 4mL of dissection media to gain an 8mL cell suspension diluted to 6% Optiprep^TM^.

*For cost reduction, PBS can be used as the diluent instead of dissection media.

17. Centrifuge 200g, 15min at RT, full brake.

*For further removal of debris, the cell pellet can be resuspended in PBS and centrifuged once more at 200g, 3min, RT full brake.

18. Resuspend the cell pellet in OPC medium, count cells, and seed the cells onto the PDL-coated culture surfaces at a seeding density of 1*10^4^/cm^2^.

*The final cell yield for one pup usually ranges between 1.5-2*10^6^, and therefore seeding at two T75 flasks per pup (=brain) will generally suffice.

***Culture maintenance, passaging, and differentiation***

19. Maintain the cultures in OPC medium. Colonies of OPCs will be visible by day 3, and 5-6 days will allow colonies to reach confluency.

*At day 3 of culture, perform a half change of OPC media, or for convenience add 30ng/mL PDGF-AA.

*Maintaining the cultures at this step longer than 6 days will cause the crowded OPCs to autodifferentiate.

20. Passage the OPCs with Accutase®. Passaged OPCs can be seeded onto different surfaces coated with PDL, at the same seeding density (1*10^4^/cm^2^)
21. Stabilize/proliferate for 2 days in OPC medium without EGF to obtain pure OPC cultures.
22. Finally, for further differentiation into mature OLs, the media should be exchanged to OL differentiation media, and maintained for 3-4 days, with a half media change at day 2.
