## Supplementary figures and images for "A primary culture method for the easy, efficient and effective acquisition of oligodendrocyte lineage cells"

### Supplementary figure 1

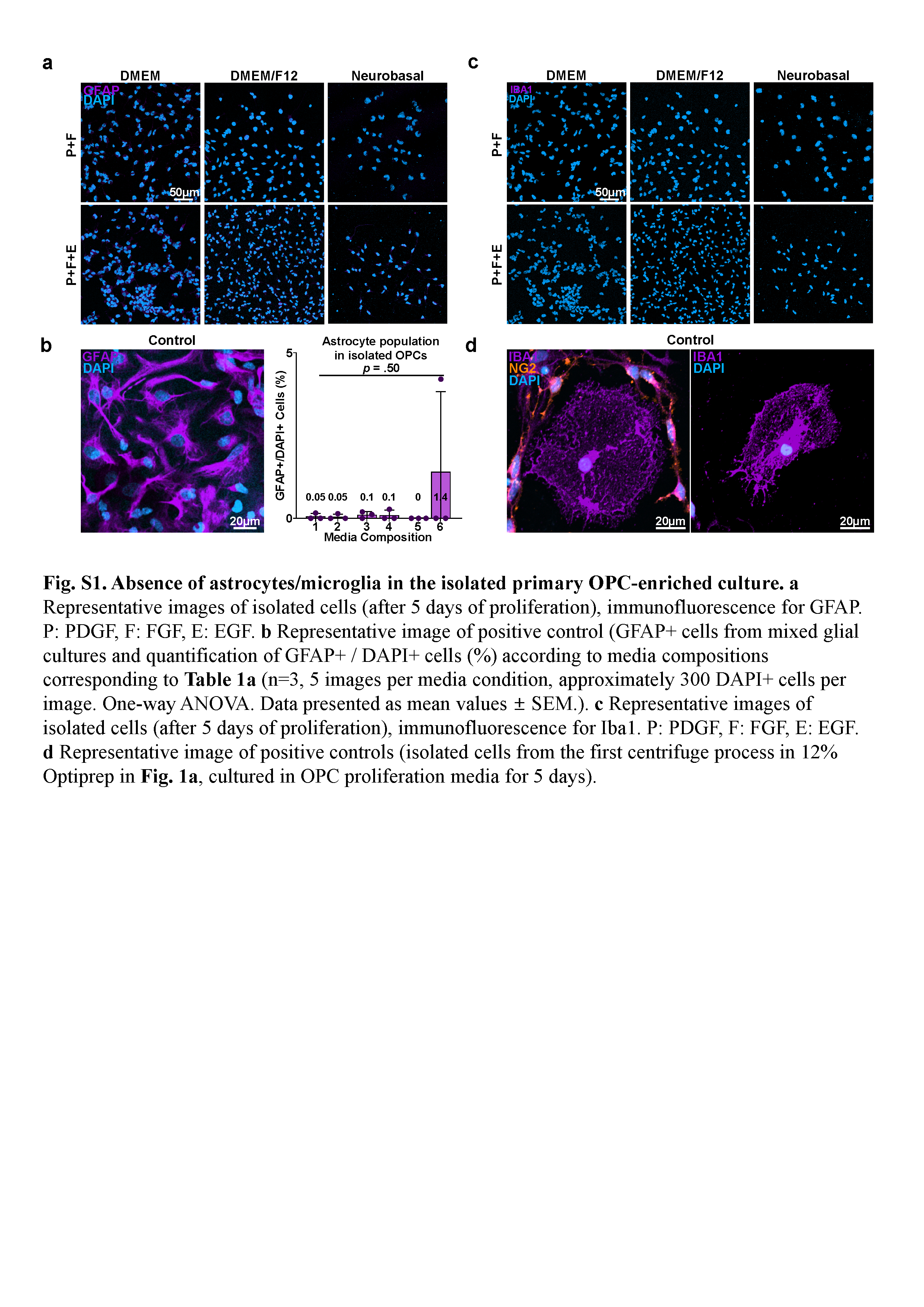

### Supplementary figure 2

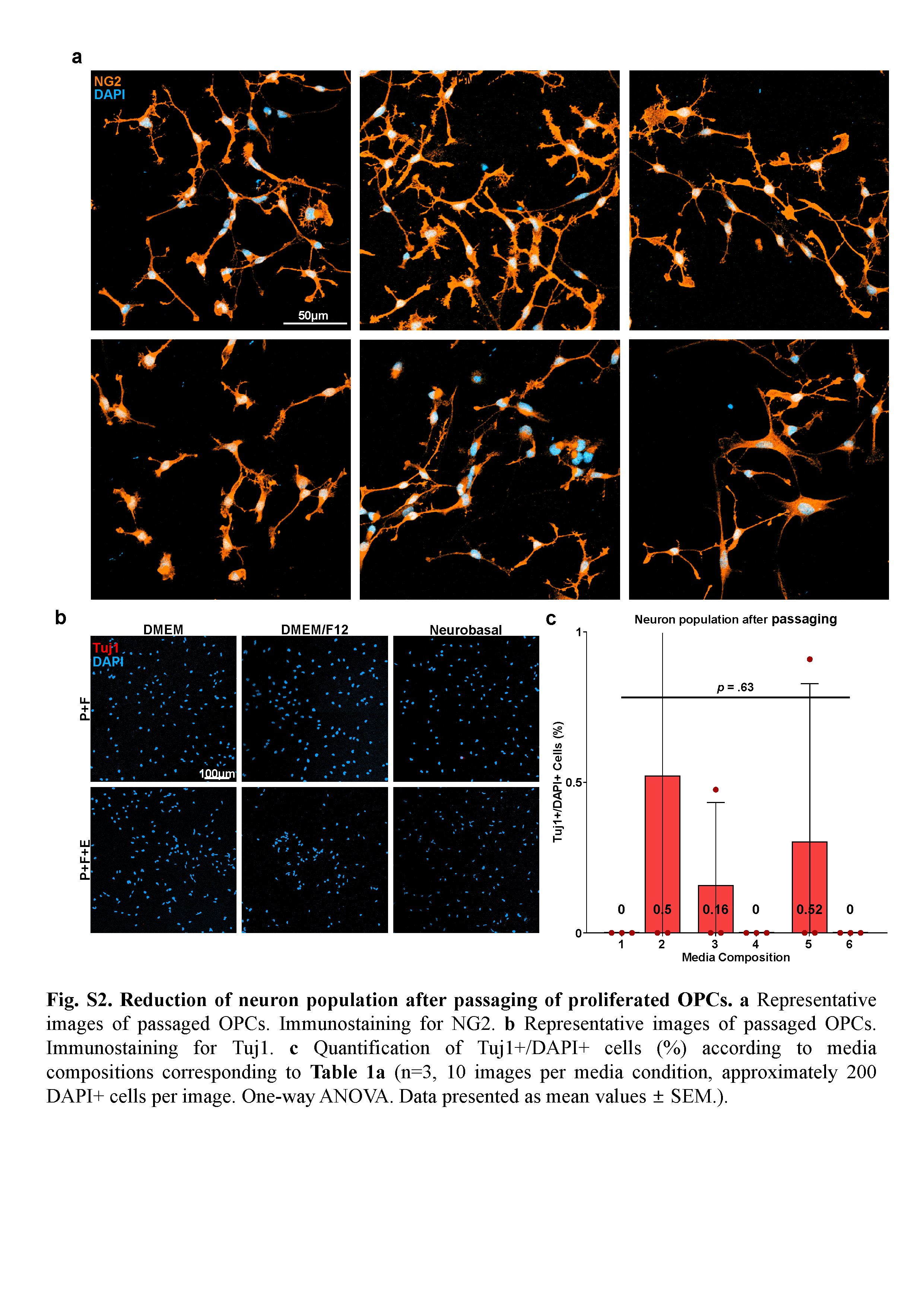

### Supplementary figure 3

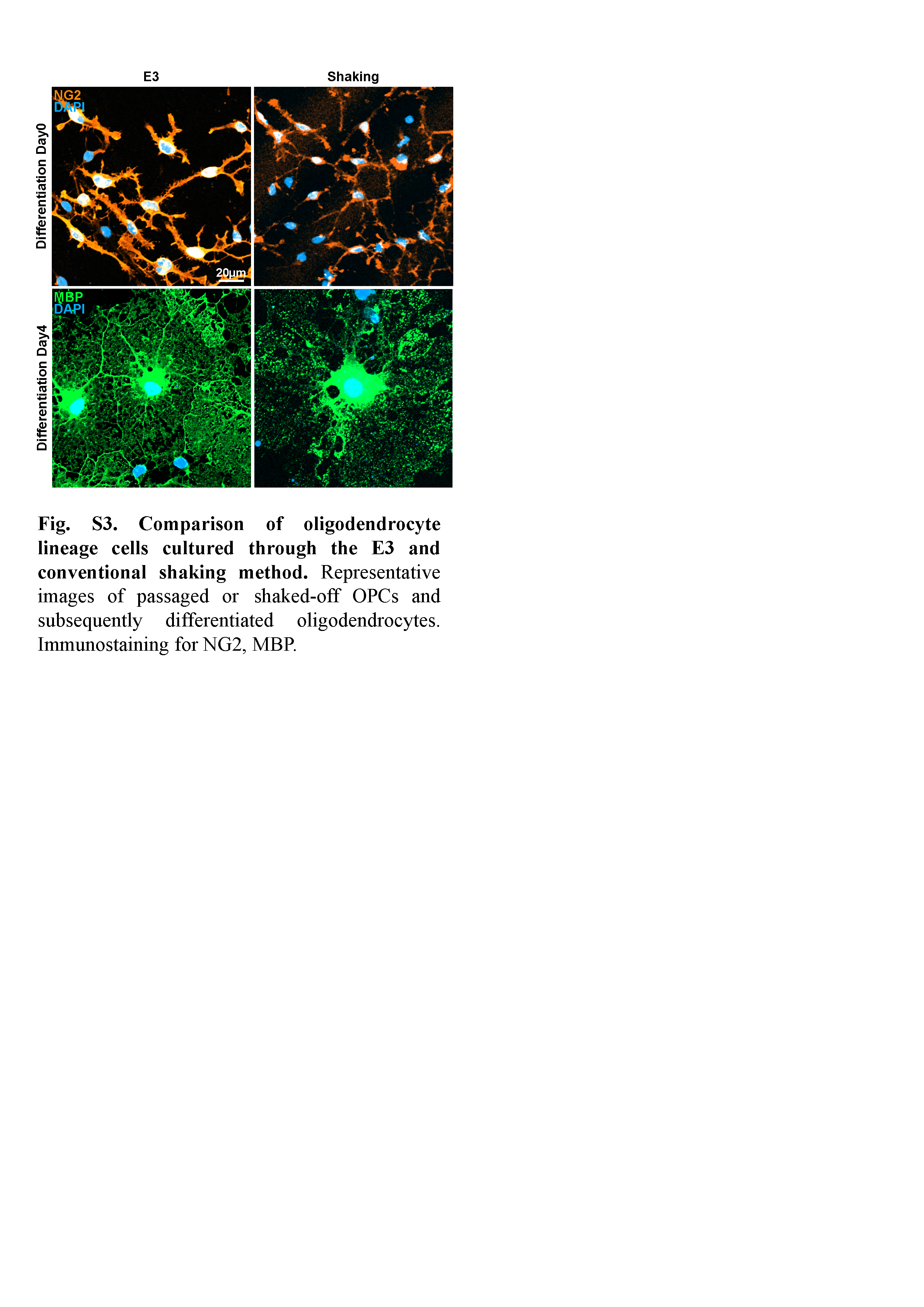
